# Impaired gephyrin G-domain trimerization and phase separation in a patient with developmental epileptic encephalopathy

**DOI:** 10.64898/2026.01.09.698633

**Authors:** Emanuel H. W. Bruckisch, Marcelo de Melo Aragão, Thais dos Santos Rhode, Ann-Kathrin Huber, Mateus de Oliveira Torres, Günter Schwarz, Filip Liebsch

## Abstract

Epilepsy, a common neurological disorder is frequently linked to genetic variants in synaptic proteins. Here, we describe a *de novo* pathogenic missense variant in the gephyrin G-domain (G134R) identified in an epileptic patient with developmental delay and seizures. Functional analyses reveal that G134R disrupts higher-order oligomerization, leading to impaired liquid-liquid phase separation (LLPS) and synaptic clustering. Recombinant G134R-gephyrin variant forms lower oligomers with reduced molybdenum cofactor (Moco) synthesis. In non-neuronal cells, G134R fails to oligomerize beyond dimers and loses Moco synthesis function. In neurons, G134R is unable to form synaptic clusters and exerts a dominant-negative effect on WT-gephyrin, severely disrupting inhibitory synapse formation. Our findings highlight a critical role for the G-domain in gephyrin self-assembly and LLPS, shifting the focus from the E-domain-centric view of gephyrin function and providing a novel molecular mechanism for epilepsy linked to G-domain mutations.

## INTRODUCTION

Epilepsy, affecting over 50 million people worldwide, is a common neurological disorder characterized by recurrent seizures caused by an imbalance between excitatory and inhibitory neuronal activity (Schwartzkroin 2012; Yardi 2025). Among the most severe forms are developmental and epileptic encephalopathies (DEE) (Scheffer et al. 2025), which often arise from *de novo* genetic mutations and manifest in early childhood with seizures and are regularly accompanied by developmental delay, and cognitive impairment (Cross and Guerrini 2013). Mutations in genes involved in synaptic transmission are frequent causes of DEE, with inhibitory synaptic proteins playing a prominent role (Guerrini et al. 2023; Uzay and Kavalali 2023; Reinthaler et al. 2015).

Gephyrin is the essential postsynaptic scaffold protein at inhibitory synapses. It interacts with glycine receptors (GlyRs) (Meyer et al. 1995) and GABA type A receptors (GABA_A_Rs) (Maric et al. 2011), thereby anchoring and clustering these receptors to maintain inhibitory strength (Fritschy et al. 2008; Tyagarajan and Fritschy 2014). Gephyrin assembles into higher-order oligomeric complexes, mediated by trimerization of its N-terminal G-domain (Schwarz et al. 2001; Saiyed et al. 2007) and dimerization of the C-terminal E-domain (Kim et al. 2006). Recently, gephyrin oligomerization has been linked to liquid-liquid phase separation (LLPS), proposed as a central mechanism for the formation and plasticity of inhibitory synapses (Bai et al. 2021; Bai and Zhang 2022; Zhu et al. 2024; Lee et al. 2024).

Besides its synaptic role, gephyrin catalyzes the final steps of molybdenum cofactor (Moco) biosynthesis, a function conserved from bacteria to humans (Belaidi and Schwarz 2013). In this pathway, the G-domain catalyzes the adenylation of molybdopterin (MPT) to generate MPT–AMP, while the E-domain subsequently hydrolyzes AMP and incorporates molybdate to yield active Moco (Belaidi and Schwarz 2013). Mutations in gephyrin can thus impair metabolic functions, leading to the fatal metabolic disorder Moco deficiency (MoCD) (Schwarz et al. 2009; Johannes et al. 2022).

Pathogenic variants and exonic microdeletions in the gephyrin gene (*GPHN*) have been identified in patients with a spectrum of neurodevelopmental and neuropsychiatric disorders, including autism spectrum disorder, schizophrenia, and epilepsy (Lionel et al. 2013). Mechanistic studies on patients demonstrated that missense variants within the E-domain, such as G375D and D422N reduce gephyrin receptor-binding and clustering, thereby disrupting inhibitory neurotransmission and causing DEE (Macha et al. 2022; Dejanovic et al. 2015). Additionally, single particle cryogenic electron microscopy analyses revealed that these two variants fail to form filaments via subdomain II interactions of adjacent dimers, which likely explains their inability to form higher oligomers and undergo LLPS (Bai et al. 2021; Kim et al. 2021; Macha et al. 2025).

Here, we describe the first pathogenic gephyrin patient variant with a single amino acid exchange located in the G-domain, derived from an epileptic patient. We identified a heterozygous *de novo* missense mutation (c.400G>C, p.Gly134Arg “G134R”) in a patient with developmental seizures born to unrelated and unaffected parents. Functional analyses revealed that G134R disrupted the ability of gephyrin to form higher-order oligomers. We hypothesize that while the E-domain serves as the primary driver of phase separation and provides the interface for receptor binding (Macha et al. 2025), the G-domain plays a crucial role in interconnecting gephyrin conjugates into higher-order oligomers, thereby enhancing the capacity for phase separation and supporting synaptic clustering. Consequently, G134R-gephyrin failed to generate functional receptor-scaffold complexes at inhibitory synapses and similar to G375D-gephyrin decreased the density of synaptic WT clusters in a dominant-negative fashion.

## RESULTS

### Clinical description of the patient with a *de novo* G134R-gephyrin variant

We report a 15-year-old female patient from Brazil, born late preterm after an uneventful pregnancy and delivery. No prenatal, perinatal, or postnatal complications were reported. Her parents are unrelated, and there is no family history of epilepsy or other neurological disorders. Developmental delay was noted within the first year of life, and at the age of nine months she was diagnosed with epilepsy. Seizures occurred predominantly during sleep and were characterized by apnea, generalized hypotonia, and episodes of tonic posturing of all four limbs followed by clonic movements lasting several minutes. During childhood, she was further diagnosed with intellectual disability and short stature.

Clinical follow-up began when the patient was 10 years old. Neurological examination at that time showed microcephaly and minor dysmorphic features, including bilateral epicanthus, upslanting almond-shaped palpebral fissures, and small, low-set ears with attached earlobes. Brain MRI performed at 5 years of age revealed no structural abnormalities. Electroencephalography (EEG) at 10 years demonstrated epileptiform discharges, predominantly in the left temporal region during drowsiness. Treatment with a benzodiazepine was initiated and resulted in good seizure control, including normalization of the EEG by age 12.

Genetic testing was performed to further investigate the cause of her condition. Both karyotyping and chromosomal microarray analysis yielded normal results. Subsequent next-generation sequencing (NGS) of a gene panel for epilepsy and metabolic disorders identified a heterozygous missense variant in the gephyrin gene (*GPHN*; NM_001024218.2: c.400G>A; p.Gly134Arg). This variant was initially classified as a variant of uncertain significance (VUS). Segregation analysis demonstrated that the variant had arisen *de novo*. The affected residue Gly134 is located within the G-domain of gephyrin (FIGURE 1A) and is conserved throughout higher eukaryotes (FIGURE 1B) suggesting an important structural role. Gly134 resides within the β-sheet 5 structure adjacent to α-helix 5, which forms the trimerization interface (FIGURE 1C). In silico prediction tools uniformly support a deleterious effect of the Gly-to-Arg exchange: PolyPhen-2 predicts the substitution to be “probably damaging” (score 1.000), SIFT predicts it to “affect protein function” (score 0.01), and MutationTaster25 classifies the variant as “deleterious”. AlphaFold3-based structural prediction suggests that the G134R substitution perturbs the β-sheet 4 element (FIGURE 1D), with the bulky, positively charged arginine side chain protruding from and disrupting the otherwise electronegative surface defined by the WT crystal structure (FIGURE 1E). Based on these findings, we decided to further study the molecular and functional consequences of G134R-gephyrin *in vitro* and in cells.

**Figure 1:**
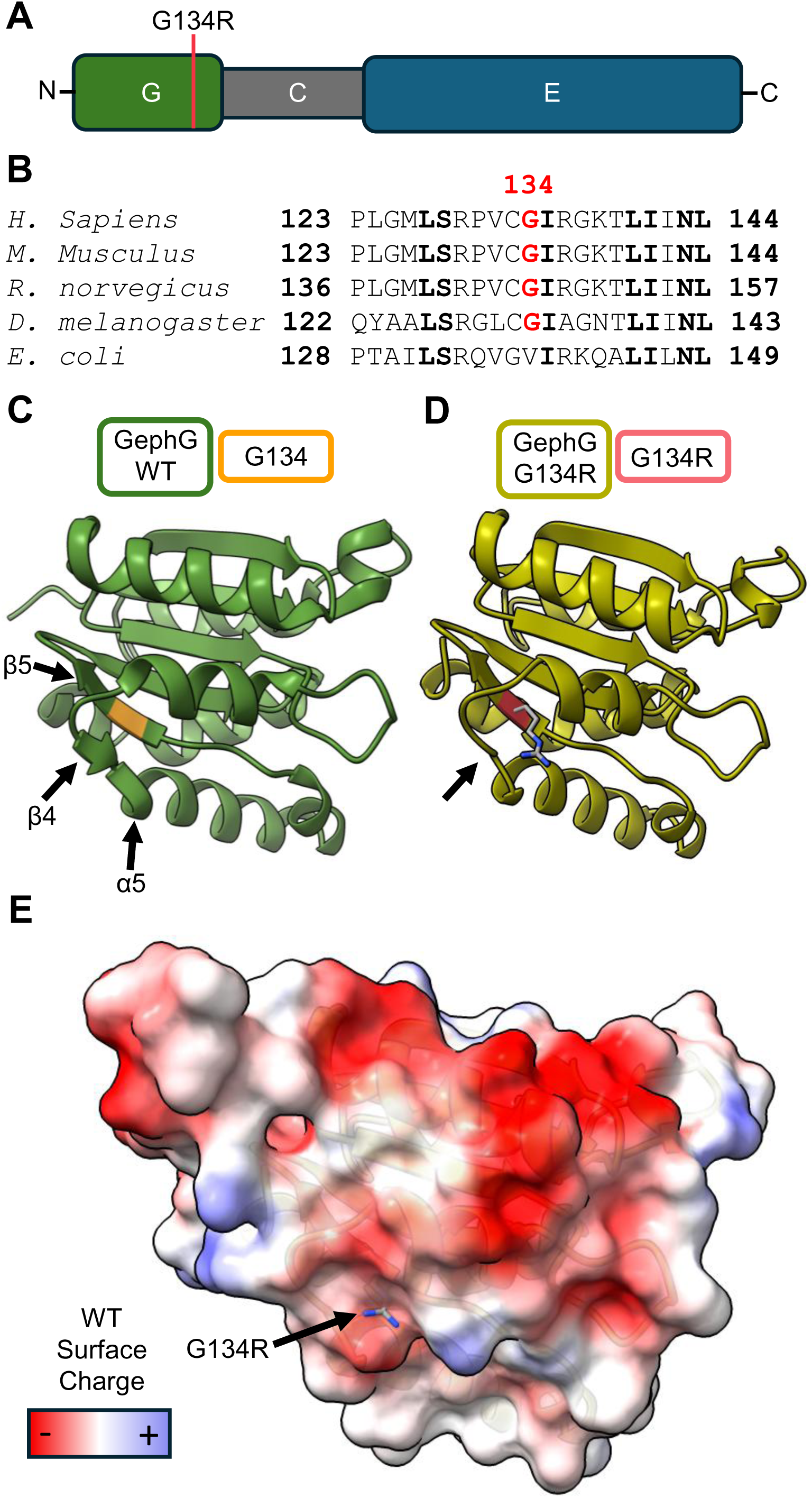
Localization of the identified *GPHN* mutation G134R. **(A)** Gephyrin domain structure with localization of G134R. **(B)** Local sequence alignment of homologous G-domains in different species. Amino acids in bold are conserved. **(C)** X-ray crystal structure of human gephyrin G-domain monomer (PDB: 1JLJ). Yellow marks glycine 134. Arrows indicate alpha-helix 5 (α5) which is important for trimerization, beta-sheet 4 (β4) and beta-sheet 5 (β5) which neighbour alpha-helix 5. **(D)** AlphaFold3 prediction of G134R-gephyrin G-domain monomer. Magenta marks the position of G134R with arginine’s carbon atoms depicted in grey and nitrogen in blue. Arrow marks the position of the lost beta-sheet 4 (β4). **(E)** Electrostatic surface prediction taken from (C) overlayed onto (D) with arrow marking G134R. Red indicates negative charge, blue positive charge. The positive arginine side chain protrudes into a net negative surface charge.

### Impaired gephyrin self-assembly and structural destabilization by G134R

To investigate the biochemical properties of G134R-gephyrin, we expressed and purified recombinant isolated G-domain variants and full-length gephyrin from *E. coli*. Size-exclusion chromatography (SEC) of the purified G-domain confirmed the expected trimeric assembly for wild-type (WT) gephyrin, yielding a single, well-defined peak at 75 mL elution volume (FIGURE 2A). In contrast, the G134R G-domain did not elute as a single trimer peak but produced partially overlapping peaks surrounding the elution of the WT peak. All fractions were positive for the G-domain as detected by Western blot analysis, indicating that the protein variant is present in multiple distinct oligomeric states (FIGURE 2B). These findings suggest that the canonical trimerization of the G-domain is altered by the G134R substitution.

**Figure 2:**
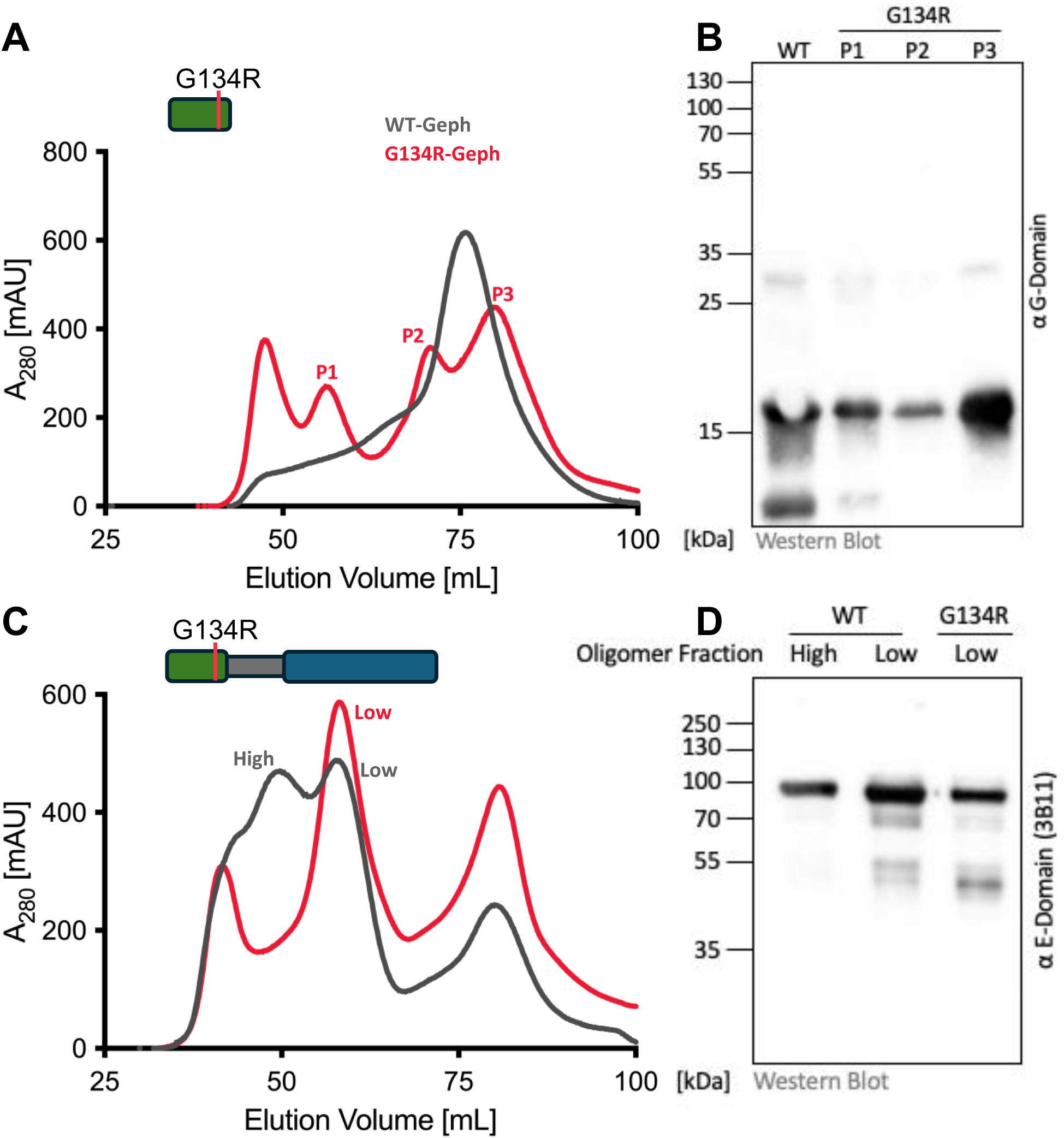
G-domain and full-length G134R-gephyrin show distinct properties *in vitro*. **(A)** Size exclusion chromatography (SEC) elution profile of G-domain WT-gephyrin (grey) and G134R-gephyrin (magenta). While WT-gephyrin shows distinct trimeric elution peak, G134R-gephyrin splits up into 4 peaks. **(B)** Western blot analysis of eluted fractions from (A) identifies G134R-gephyrin peaks P1-3 via G-domain specific antibody. The peak before 50 mL depicts cut-off peak at void volume. **(C)** SEC elution profile of full-length WT-gephyrin (grey) and G134R-gephyrin (magenta). While WT-gephyrin shows elution peaks for trimeric low-oligomerization gephyrin and an additional peak for high-oligomerization, G134R-gephyrin is missing the latter. Peaks before 50 mL and after 75 mL depict cut-off peaks at void volumes. **(D)** Western blot analysis of eluted fractions from (C) identifies G134R-gephyrin via E-domain specific antibody (3B11). Only low-oligomer fractions show characteristic double band behavior.

Analysis of full-length gephyrin revealed the characteristic oligomerization elution-profile previously described for WT protein, with specific populations corresponding to lower trimeric and higher oligomeric assemblies (50 mL and 60 mL respectively in FIGURE 2C), in addition to aggregates eluting at 40 mL and degradation products eluting at 80 mL. Strikingly, full-length G134R-gephyrin completely lacked the high-oligomer peak at 50 mL, while only the low trimeric fraction was detectable. These trimers eluted at 60 mL, migrated at ∼85 kDa on SDS–PAGE, and were indistinguishable from WT gephyrin by Western blot analysis, indicating that these units remain intact but fail to engage in higher-order assembly (FIGURE 2D).

Given these profound alterations in the oligomerization properties of G134R-gephyrin, we next asked whether the protein retains its biological functions. We therefore performed in-depth biochemical analyses of the two major activities of gephyrin: clustering of inhibitory neurotransmitter receptors and catalysis of the molybdenum cofactor (Moco) biosynthesis.

### G134R-gephyrin showed unaffected receptor binding and reduced but functional Moco synthesis activity

E-domain dimers of gephyrin mediate the postsynaptic anchoring of inhibitory receptors by binding to conserved intracellular loop domains of GlyR β- and GABA_A_R α3-subunits. Although Gly134 is located distantly in the N-terminal G-domain, the loss of higher-order oligomerization in G134R-gephyrin prompted us to test whether this variant affects receptor binding. To this end, we performed isothermal titration calorimetry (ITC) experiments using a synthetic peptide derived from the GlyR β intracellular loop domain (GlyR β-ICD). As previously reported (Macha et al. 2022), this peptide is a suitable model to study gephyrin–receptor interactions since GlyRs and GABA_A_Rs compete for the same binding site on gephyrin (Maric et al. 2014). In agreement with earlier studies, WT-gephyrin showed two distinct binding sites, a high-affinity interaction in the sub-micromolar range (K*_D_* = 0.27 µM, N = 0.33) and a low-affinity interaction in the double-digit micromolar range (K*_D_* = 15.11 µM, N = 0.71) (FIGURE 3B). Strikingly, the G134R variant displayed binding curves and thermodynamic parameters highly similar to those of WT protein (FIGURE 3A), with no statistically significant differences in affinity or stoichiometry (FIGURE 3B). These results indicate that, despite its profound oligomerization defects, G134R-gephyrin retains intact receptor-binding capacity *in vitro*.

**Figure 3:**
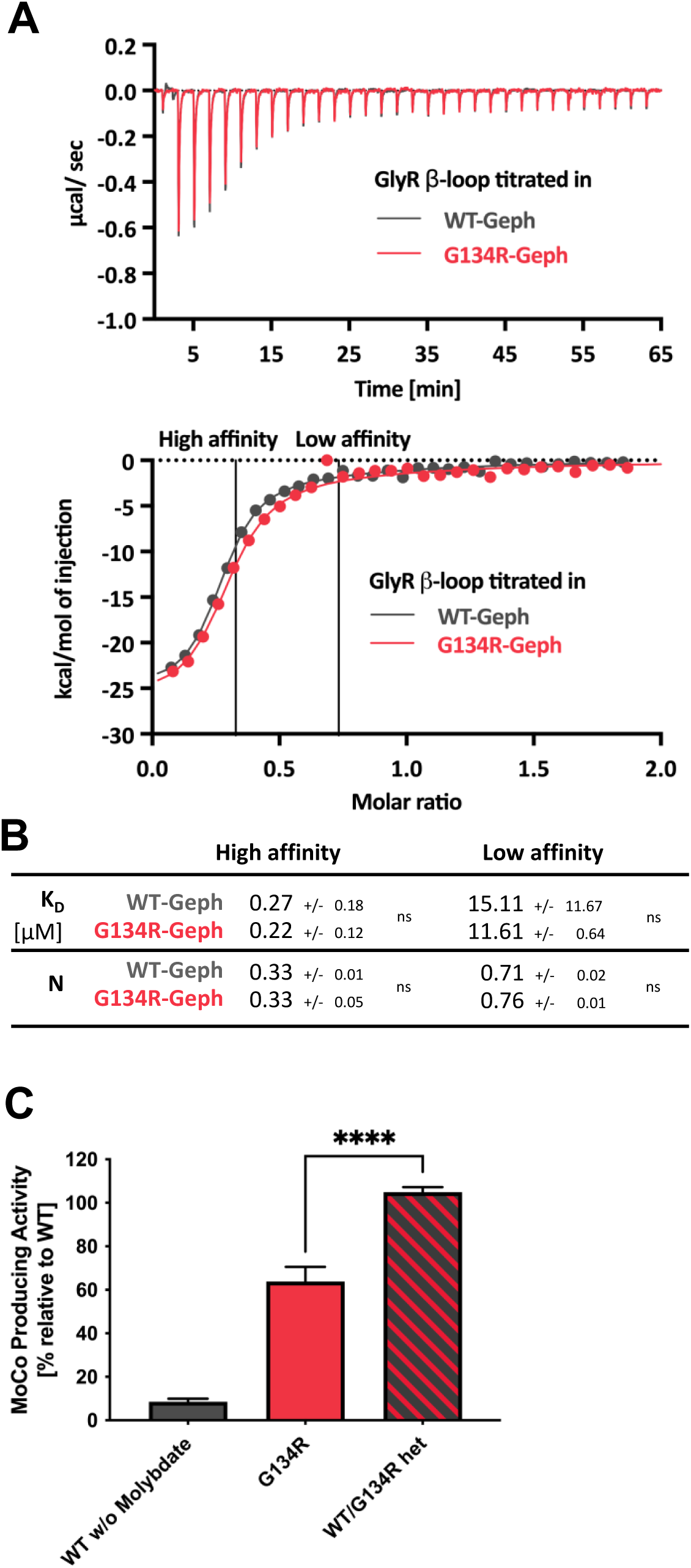
Preserved GlyR β-Loop binding and Moco activity in G134R-gephyrin *in vitro*. **(A)** Representative isothermal titration calorimetry (ITC) binding curves illustrating the interaction of the GlyR β-loop with either WT-gephyrin or G134R-gephyrin are shown. Experimental data were fitted with a two-site binding model (solid lines), distinguishing high-and low-affinity interaction sites. **(B)** Quantitative binding parameters obtained from the ITC measurements, including dissociation constants (KD, μM) and stoichiometry values (N), are presented. Data are reported as mean ± SD (n = 3, derived from three independent protein purifications) and statistical comparisons were performed using an unpaired, two-tailed Student’s t-test. No significant differences were detected for any of the parameters (ns, p ≥ 0.05). **(C)** In vitro molybdenum cofactor (Moco) biosynthesis assay was performed using 150 pmol of purified 6His-tagged gephyrin proteins as described earlier (Belaidi and Schwarz 2013). Presented as % relative to WT-gephyrin activity. Reaction lacking molybdenum (w/o molybdate) were included as internal negative control to validate assay performance. Data are reported as mean ± SD (n = 3, derived from three independent measurements) and statistical comparisons were performed using an one-way ANOVA (F = 170.5, p < 0.0001) with Tukey’s multiple comparisons test (p_G134R vs WT/G134R het_ < 0.0001).

Gephyrin is a moonlighting protein with a second essential function in cellular metabolism: catalysis of the final two steps in Moco biosynthesis. We therefore assessed whether structural alterations in the G-domain of G134R-gephyrin impair its enzymatic activity. *In vitro* Moco synthesis assays revealed that G134R-gephyrin variant produced approx. 400 pmol Moco, corresponding to ∼50 % of WT activity (FIGURE 3C). Thus, although the G134R substitution markedly reduces catalytic efficiency, it does not completely abolish enzymatic activity. Importantly, when WT and G134R proteins were mixed in a 1:1 ratio mimicking the heterozygous expression condition in the patient, Moco production was restored to ∼700 pmol, being indistinguishable from WT (FIGURE 3C). These findings suggest that G134R-gephyrin does not exert a dominant-negative effect on the enzymatic function of WT-gephyrin.

Taken together, our results demonstrate that G134R-gephyrin preserves its two core functions *in vitro*: receptor binding and Moco synthesis. The partial Moco activity is likely mediated by the trimers that still form *in vitro* and are presumed to remain catalytically competent. Consistent with the absence of structural brain abnormalities on MRI and with the lack of biochemical or clinical signs of Moco deficiency this is not expected to be clinically relevant. Instead, our data point toward defective higher-order self-assembly as the critical impairment of this variant. Given the established link between gephyrin receptor clustering and LLPS at inhibitory synapses, we next investigated whether the inability of G134R-gephyrin to form higher-order assemblies translates into impaired phase separation capacity.

### Loss of higher-order oligomers impairs gephyrin phase separation

Given that gephyrin synaptic clustering has been linked to LLPS at inhibitory synapses, we next tested whether the inability of gephyrin-G134R to form higher oligomers alters its phase separation behavior. We employed an established assay (Bai et al. 2021), in which the ICDs of inhibitory receptors are fused to the dimeric yeast zinc finger protein GCN4 and used them to probe receptor–induced gephyrin phase separation in sedimentation experiments (FIGURE 4A). As described previously, the GlyR β-ICD represents the strongest known gephyrin interactor, the GABA_A_R α3-ICD shows intermediate affinity, and the GABA_A_R α1-ICD interacts only weakly.

**Figure 4:**
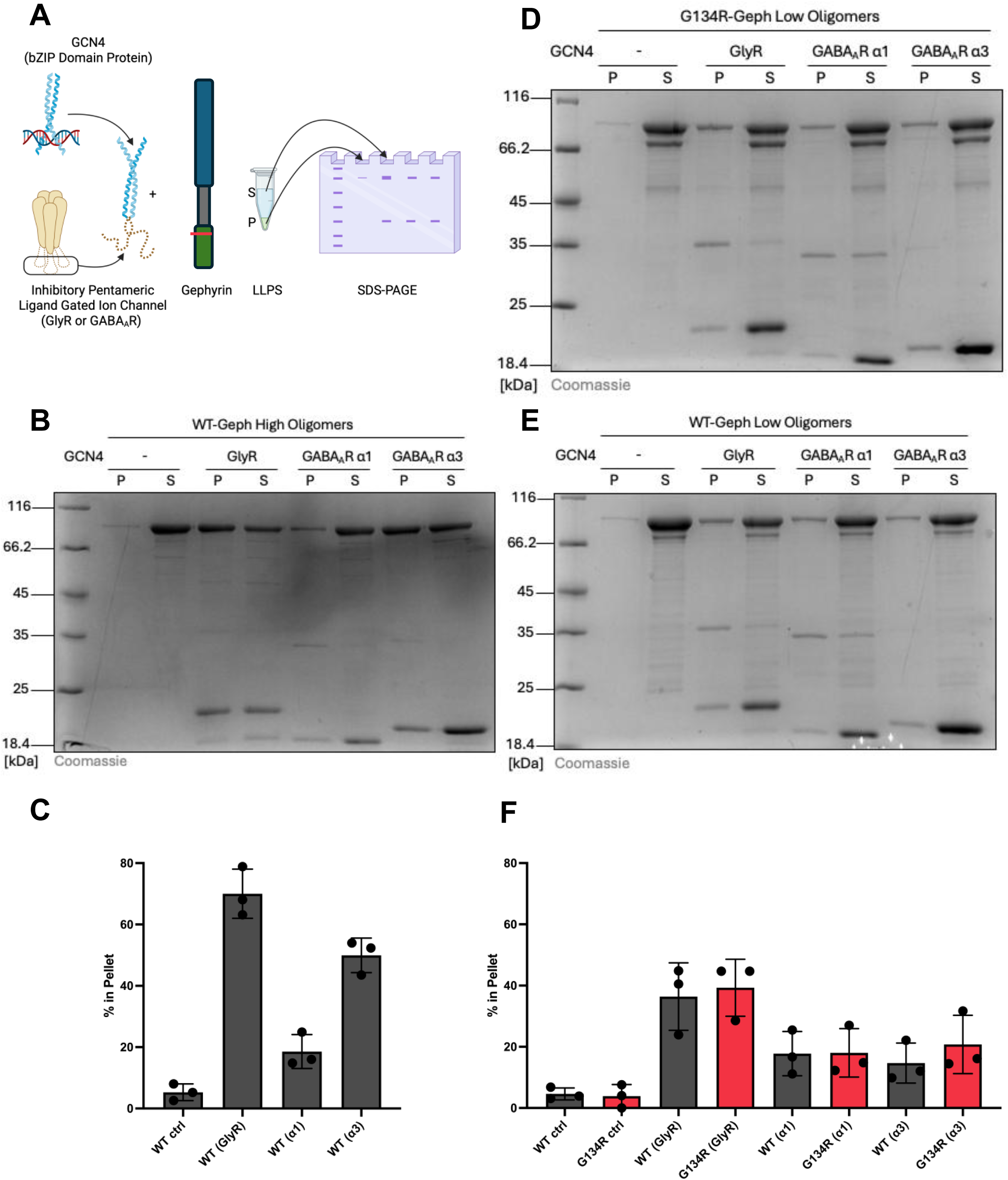
A loss of higher oligomers reduces G134R gephyrin LLPS *in vitro*. **(A)** Cartoon depicting GCN4-based receptor models (Bai et al. 2021) and experimental design of liquid-liquid phase separation (LLPS) sedimentation assay. **(B, D, E)** Representative Coomassie stained SDS-PAGE of LLPS sedimentation assay from WT-gephyrin high oligomers (B), G134R-gephyrin low oligomers (D) and WT-gephyrin low oligomers (E) with GCN4-based receptor models for GlyR, GABA_A_R-α1 and GABA_A_R-α3 showing the amount of protein present in pellet (P) and supernatant (S) fractions. **(C, F)** The band intensities in the pellet fraction were measured and are shown as mean ± SD from three independent experiments (n = 3). Panel C is based on the data in (B), while panel F summarizes the data from (D) and (E).

When tested with the high oligomeric fraction of WT-gephyrin (isolated after SEC purification), we observed robust receptor-specific phase separation. Approximately 70 % of gephyrin were recovered in the sedimented pellet fraction in presence of the GlyR β-ICD fusion protein, ∼50 % with GABA_A_R α3-ICD, and ∼20 % with GABA_A_R α1-ICD, consistent with the expected order of interaction strengths (FIGURE 4B and C). In contrast, the lower trimeric fraction of WT gephyrin showed markedly reduced phase separation, with GlyR β-ICD and α3-ICD pelleting decreased to ∼40 % and ∼20 %, respectively, while α1-ICD interactions remained unchanged (FIGURE 4E and F). These results highlight the critical contribution of higher-order oligomers for phase separation. G134R-gephyrin, which lacks the ability to form higher oligomers, behaved similarly to WT trimers in LLPS (FIGURE 4D and F). However, since the variant protein lacks higher-order assemblies, the majority of the phase separation–competent protein population was lost. We therefore conclude that G134R-gephyrin exhibited a strongly reduced propensity for LLPS, due to its inability to form higher oligomers.

Importantly, these findings suggest that although receptor binding to the E-domain is preserved, the structural instability of gephyrin assemblies lacking higher oligomers may compromise the integrity of the inhibitory postsynaptic density assuming that G134R-gephyrin fails to generate the protein-dense, phase-separated scaffold required for stabilizing receptor clusters, thereby weakening inhibitory inputs. To directly test this hypothesis in a physiological setting, we next transitioned from *in vitro* assays to cellular models, first expressing G134R-gephyrin in HEK cells lacking endogenous gephyrin expression to assess both its oligomeric state, Moco forming activity and whether its altered LLPS propensity manifests as discrete fluorescent condensates in living cells. Building on these experiments, we then extended our analysis to primary hippocampal neurons to determine whether G134R-gephyrin can still support the formation of functional inhibitory synapses.

### In HEK cells, G134R-gephyrin fails to oligomerize, resulting in disrupted Moco synthesis and absence of LLPS

Our HEK cell-based experiments reveal a severe defect in oligomerization of the G134R-gephyrin variant compared to both WT-gephyrin and previously characterized control variants. Using blue native PAGE, we demonstrate that G134R fails to form native trimers or higher oligomers, running at the same molecular weight as the dimeric control (G2-gephyrin splice variant; (Saiyed et al. 2007; Bedet et al. 2006)), while WT-gephyrin migrates above the trimeric control (S505F-gephyrin E-domain mutant; (Burdina et al. 2025)), confirming its ability to assemble into larger complexes (FIGURE 5A). This defect is more pronounced in cells than *in vitro*, where trimers were still observed. To assess functional consequences, we co-expressed WT- or G134R-gephyrin with sulfite oxidase (SUOX) in HEK gephyrin knockout cells (KO). SUOX activity, which depends on Moco biosynthesis by gephyrin, was restored only by WT-gephyrin, indicating that G134R lacks sufficient catalytic activity for Moco synthesis in this context (FIGURE 5B). Similar protein expression was validated via Western blot analysis (FIGURE 5C). Fluorescence microscopy revealed that overexpressed mScarlet-tagged WT-gephyrin forms discrete cytoplasmic condensates, consistent with LLPS (Lee et al. 2024), while G134R remains diffusely distributed (FIGURE 5D). Co-expression with the mEGFP-tagged dimeric receptor model GCN4-GlyR further enhanced WT LLPS but failed to induce condensate formation by G134R (FIGURE 5E). These findings indicate that the G134R variant disrupts both oligomerization and phase separation in cells, impairing gephyrin’s metabolic and scaffolding functions. Building on these results, we next aimed to extend our analysis to neuronal cell culture to investigate whether G134R-gephyrin can still support the formation of functional inhibitory synaptic clusters.

**Figure 5:**
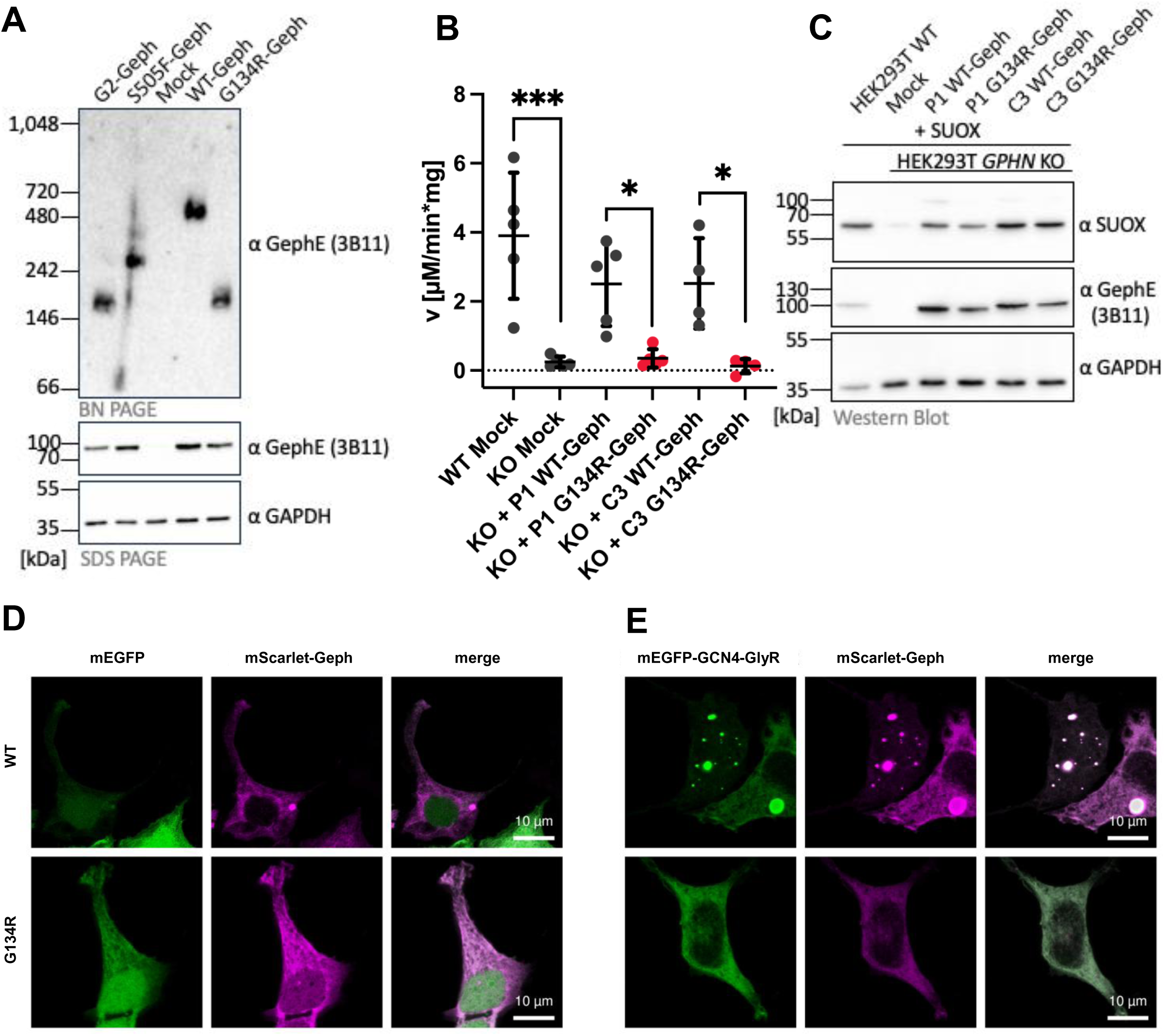
Loss of oligomerization in G134R-gephyrin abolishes Moco synthesis and LLPS in HEK cells. **(A)** Representative Western blot analysis after blue native PAGE of HEK293T *GPHN* KO cells transfected with indicated gephyrin variants. As control Western blot analysis of SDS-PAGE was performed. **(B)** Sulfite oxidase-specific activity in whole cell lysates. Either HEK293T WT (WT) or *GPHN* KO (KO) were transfected with indicated gephyrin variants or empty plasmid (Mock) (n = 5 independent experiments; mean ± SD are indicated). Data were analyzed by one-way ANOVA (F_5,22_ = 10.524, p = 2.94 * 10^-5^), Tukey’s post hoc test; p_WT Mock vs. KO Mock_ = 0.000241; p_KO + P1 WT-Geph vs. KO + P1 G134R-Geph_ = 0.0416; p_KO + C3 WT-Geph/KO vs. C3 G134R-Geph_ = 0.0435 **(C)** Representative Western blot analysis of cell lysates from (B). **(D, E)** Representative confocal microscopy images of HEK *GPHN* KO cells expressing mScarlet tagged WT-gephyrin or G134R-gephyrin in the presence of mEGFP (D) or mEGFP tagged GCN4-GlyR (E). Scale bars represent 10 μm.

### G134R-gephyrin disrupts neuronal clustering and acts dominant negatively when co-expressed with WT-gephyrin

We finally tested the G134R variant in primary hippocampal neurons. To achieve expression in cells lacking endogenous gephyrin, we generated dissociated hippocampal cultures from floxed gephyrin mice (O’Sullivan et al. 2016). Knockout of endogenous gephyrin was achieved by Cre expression, which was combined with Cre-dependent expression of mScarlet-tagged WT- or G134R-gephyrin (FIGURE 6A). Under these conditions, WT-gephyrin formed robust somatic and dendritic clusters colocalizing with the presynaptic inhibitory marker vGAT and the γ2 subunit of GABA_A_R, confirming their synaptic identity (FIGURE 6B and C). In contrast, neurons expressing only G134R-gephyrin displayed a dramatic loss in the number of synaptic clusters, with markedly fewer puncta colocalizing with vGAT and γ2 (FIGURE 6B and C). These results demonstrate that G134R-gephyrin is unable to fully support the assembly of inhibitory postsynaptic scaffolds in neurons by itself.

**Figure 6:**
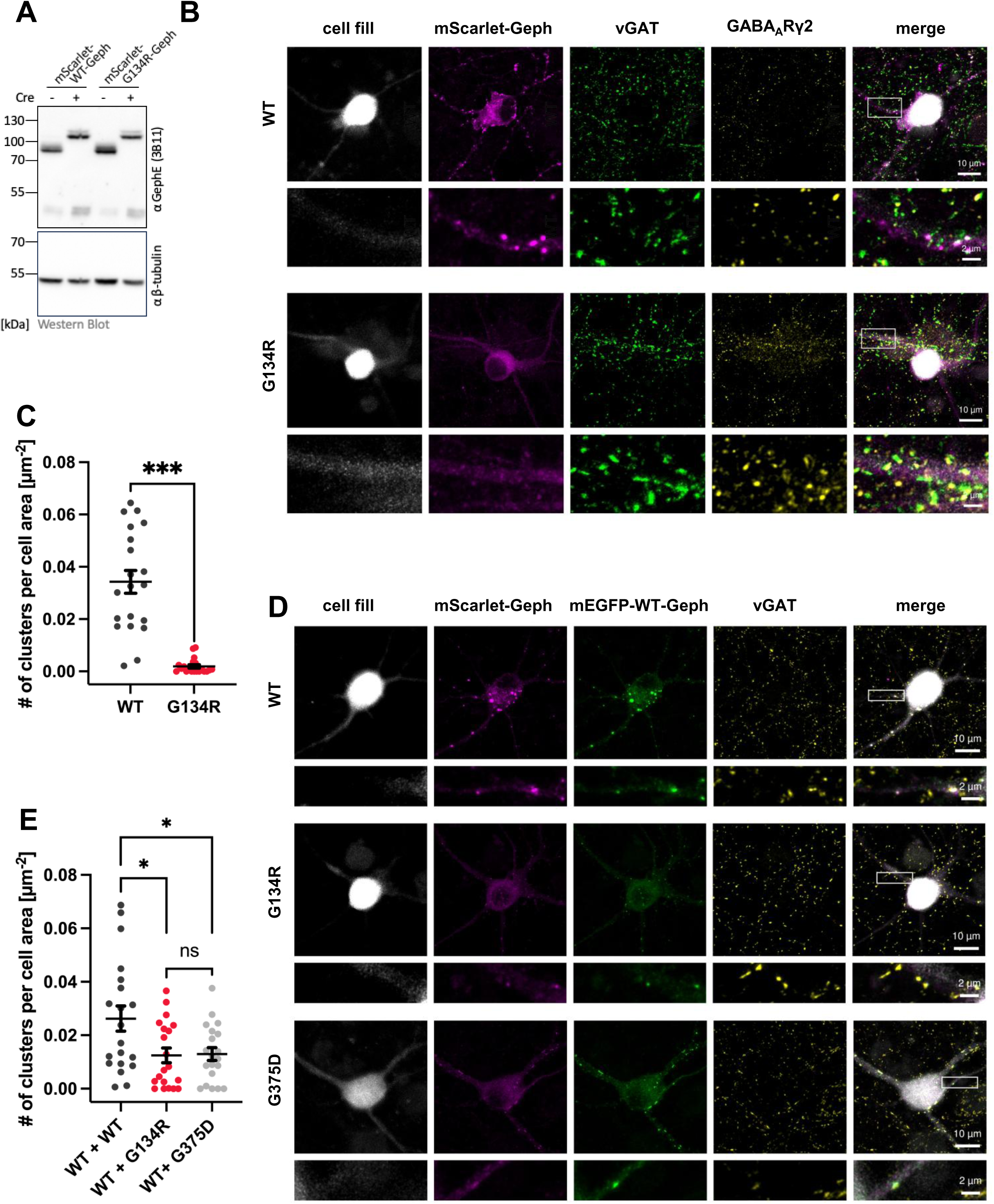
Differential expression of WT- and G134R-gephyrin in primary hippocampal neuron culture. **(A)** Representative Western blot analysis of lysates from gephyrin floxed/floxed murine hippocampal neurons infected with moxBFP (-) or moxBFP-P2A-Cre (+) together with either mScarlet-WT-Geph or mScarlet-G134R-Geph. **(B, D)** Representative confocal images of gephyrin^floxed/floxed^ murine hippocampal neurons transfected with moxBFP-P2A-Cre and (B) infected with either mScarlet-WT-Geph or mScarlet-G134R-Geph and immunostained against vesicular GABA transporter (vGAT, green) and GABA_A_R γ2 subunit (yellow) or (D) infected with mEGFP-WT-Geph and either mScarlet-WT-Geph or mScarlet-G134R-Geph and immunostained against vesicular GABA transporter (vGAT, yellow). **(C, E)** Quantification of mScarlet-tagged gephyrin clusters was carried out using automated synapse analyses. Each data point represents synaptic cluster density (# per cell area µm^-2^) per acquired image; n = 20 cells per condition from five independent cultures, mean ± SD are indicated. Data were analyzed by (C) Mann-Whitney U test U = 391, p = 0.00000023, effect size r = 0.820 (E) one-way ANOVA (F_2,57_ = 5.139, p = 0.009), Tukey’s post hoc test; p_WT + WT vs. WT + G134R_ = 0.0176; p_WT + WT vs. WT + G375D_ = 0.0228; p_WT + G134R vs. WT + G375D_ = 0.995

To model the heterozygous state found in the patient, we tested whether the G134R variant exerts a dominant-negative effect on WT-gephyrin, potentially contributing to the observed epileptic phenotype. We co-expressed mEGFP-tagged WT-gephyrin and mScarlet-tagged G134R in neurons, enabling clear distinction between the two proteins based on their respective fluorescent labels. As a positive control, we included the previously described dominant-negative variant G375D (FIGURE 6D). Heterozygous expression of G134R significantly reduced the density of WT inhibitory postsynaptic clusters to a similar extent as G375D, supporting a dominant-negative effect (FIGURE 6E). These findings indicate that G134R-gephyrin interferes with WT-gephyrin function, providing a plausible molecular mechanism for the patient’s epileptic phenotype.

## DISCUSSION

Our study identifies G134R-gephyrin as the first epilepsy patient-derived variant from a missense mutation in the *GPHN* gene affecting the G-domain of gephyrin, demonstrating that a single amino acid substitution disrupts gephyrin’s oligomerization and LLPS. Previously, two missense mutations affecting gephyrin’s G-domain, namely V43L and A91T were identified in patients with autism spectrum disorder (Lionel et al. 2013). However, functional analyses revealed only moderate clustering deficiencies for A91T, while V43L behaved normally under the experimental conditions (Kim et al. 2021). Recombinant and purified G134R-gephyrin formed trimeric lower-order oligomers, shows reduced Moco synthesis, while a dominant-negative effect on its enzymatic activity was not observed as expected from the patient’s biomarker profile. However, in a human cellular model, G134R-gephyrin failed to form higher-order oligomers beyond E-domain mediated dimerization and lost its Moco synthesis function entirely. While G134R-gephyrin retains the ability to interact with the intracellular domain of the glycine receptor, it is unable to form synaptic clusters in dissociated hippocampal neurons, both in a homozygous and heterozygous context, where it acts dominantly negative on WT-gephyrin. This dominant-negative effect, which mirrors the behavior of the epileptogenic E-domain mutant G375D (Dejanovic et al. 2015), provides a direct molecular explanation for the patient’s epileptic phenotype by effectively depleting functional inhibitory receptor-scaffold complexes at synapses.

Previous work from our laboratory identified the two E-domain gephyrin variants G375D and D422N in patients with developmental epilepsy. Both variants perturb LLPS, thus establishing the E-domain as a crucial structural hub for condensate formation and inhibitory synapse integrity (Dejanovic et al. 2015; Macha et al. 2022; 2025). Structural and functional analyses revealed that the subdomain two (SDII) within the E-domain is essential for the recently discovered filament formation via SDII–SDII interactions, which in turn underlie LLPS and define the structural basis of postsynaptic gephyrin scaffolds (Macha et al. 2025). The G375D variant, located within SDII, causes unfolding of this subdomain, abolishing SDII–SDII mediated filament formation and thereby eliminating LLPS and synaptic clustering capacity (Macha et al. 2025). In contrast, the D422N variant disrupts a critical salt bridge with R379, yet preserves the overall SDII fold, still resulting in loss of LLPS and impaired inhibitory synaptic transmission, highlighting that subtle perturbations of the SDII network suffice to destabilize condensate formation.

Taken together with our current observations, we propose a refined model for establishing the postsynaptic architecture: while the E-domain is the primary driver of phase separation and provides the receptor-binding interface, the G-domain is crucial for interconnecting E-domain gephyrin conjugates into higher-order oligomers, thereby amplifying LLPS capacity and stabilizing synaptic clustering. The strong G-domain trimerization (Herweg and Schwarz 2012) may provide a hierarchical organizing principle that ensures E-domain dimers efficiently find and engage each other, thereby reaching the threshold concentrations required for LLPS and filament maturation under physiological conditions. Simultaneously, E-domain dimers nucleate filaments through SDII–SDII interactions, and G-domain trimerization could interlink these E-domain-based filaments into a robust three-dimensional scaffold that can maintain the protein concentrations and geometric connectivity observed *in vitro* by cryo-EM and required *in vivo* at synapses. Within this framework, G134R-gephyrin is as functionally disruptive at the cellular and clinical levels as the E-domain SDII variant G375D, despite residing outside the critical receptor binding and filament forming domains of gephyrin, indicating that G-domain-mediated higher-order oligomerization is equally important as SDII-dependent filament assembly for LLPS and synapse formation.

Our convergent mechanistic insight suggests an additional evolutionary advantage for the fusion of G- and E-domains via the flexible C-domain in vertebrates within a single polypeptide chain: it serves both, product-substrate-channeling in Moco synthesis (Belaidi and Schwarz 2013) and precise control over condensate assembly for postsynaptic receptor clustering. The flexible C-domain likely permits the G-domain trimeric core to be spatially separated from the E-domain filament lattice, enabling dynamic regulation of scaffold connectivity without steric interference with filament formation.

The central C-domain of gephyrin, which behaves as an intrinsically disordered region (IDR), is indispensable for scaffold condensation and can be functionally replaced by heterologous IDRs, in some cases even enhancing gephyrin’s LLPS capacity, indicating that the C-domain acts as a charged spacer that thermodynamically modulates phase separation rather than as a classical self-associating IDR (Lee et al. 2024). In apparent contrast, segments of the C-domain can inhibit LLPS driven by the E-domain *in vitro*, suggesting that under these conditions C-domain–E-domain contacts can restrain E-domain phase separation and thus provide a built-in brake on spontaneous condensation (Bai et al. 2021). Integrating these findings, the conformation and exposure of the C-domain operates as a dynamic hub that senses local receptor density and postsynaptic architecture, defining the boundary of the inhibitory postsynaptic density so that gephyrin condensates remain confined to receptor-enriched nanodomains, while remaining permissive to rapid remodeling. In this scenario, the G-domain not only interconnects E-domain filaments but also allosterically controls C-domain compaction *versus* extension, thereby toggling between C-domain–shielded states that limit LLPS and C-domain–exposed states in which favorable electrostatic C–E interactions and increased effective valency promote percolation of a gephyrin network. This polydisperse nature of gephyrin conformation has been seen already in early structural studies using single angle X-ray spectroscopy (Sander et al. 2013). From the perspective of percolation-based phase separation hypothesis (Mittag and Pappu 2022; Shen et al. 2023), where the condensed phase emerges once a system-spanning, multivalent interaction network forms, the G-domain-mediated trimerization becomes a fundamental requirement: by providing strong, high-valency crosslinks between SDII-based E-domain filaments, G-domain connectivity helps to drive the system across the percolation threshold into a macroscopically connected, viscoelastic condensate, as described for other synaptic and PSD-like assemblies (Zhu et al. 2024). Consequently, pathogenic disruption of G-domain trimerization, as here discovered for G134R-gephyrin, is predicted not only to lower LLPS propensity but to prevent formation of a percolated scaffold network that is necessary to immobilize and concentrate inhibitory receptors within the postsynaptic density, thereby unifying G-domain and E-domain mutations within a common percolation framework for gephyrin-dependent inhibitory synapse failure.

The observation that both E-domain (G375D, D422N) and G-domain (G134R) epilepsy variants converge on a shared deficit in gephyrin LLPS implies that fine-tuning of G-domain trimerization and E-domain filament assembly constitutes a key regulatory axis to strengthen or weaken inhibitory synaptic inputs. Such multilayered regulation of filament formation, trimer crosslinking, and LLPS likely equips gephyrin with the ideal biophysical properties to function as a tunable postsynaptic scaffold in neurons, while at the same time making the system exquisitely sensitive to disease-causing missense perturbations in both the G- and E-domains.

## MATERIAL AND METHODS

### Expression constructs

6His-tagged gephyrin and GlyR β-loop-intein expression vectors have been described before (Macha et al. 2022; Dejanovic et al. 2015). GCN4 based GlyR-ICD and GABA_A_R-ɑ1/3-ICD were kindly provided by the Zhang lab (Bai et al. 2021). 6His-tagged gephyrin G-domain and full-length gephyrin (isoforms P1 and C3 in pQE80) G134R variants were cloned by site-directed mutagenesis using forward (GACCAGTGTGTAGAATAAGAGGGAAAA) and reverse (TTTTCCCTCTTATTCTACACACTGGTC) primers. 6His-tagged full-length gephyrin (isoforms P1 in pQE80) S505F variant was cloned by site-directed mutagenesis using forward (CTCATTCCCTGTGAACATAACGGCAAC) and reverse (GTTGCCGTTATGTTCACAGGGAATGAG) primers. 6His-tagged full-length gephyrin G2 in pQE80 was cloned by overlap extension PCR using G2 forward (CACGAGATGTCACTCCAGAGAAATTCCCAACATTCCCATTTTGTGGGC) and G2 reverse (CCCGTTCTATTACTTCTTTTGTGGCCCCTTTCTGGAGCCCACAAAATGGGAAT) with gephyrin flanking forward (GACCGAGGGAATGATCCT) and flanking reverse (AGGATCATTCCCTCGGTC) primers. Full-length gephyrin and its variants were subcloned into pcDNA™3.1/Zeo (+) (Invitrogen) using NEBuilder HiFi DNA Assembly (NEB). Gephyrin was initially subcloned into pmEGFP-C2 (Liebsch et al. 2023), derived from pEGFP-C2 (Clonetech) using XhoI and KpnI restriction sites. mScarlet-Gphn constructs in this plamid were generated by replacing EGFP with mScarlet from pAAV-hSyn-mScarlet (a gift from Karl Deisseroth, Addgene plasmid #131001; http://n2t.net/addgene:131001; RRID:Addgene_131001) using NEBuilder HiFi DNA Assembly (NEB). mEGFP-GCN4-GlyR in pEGFP-C2 (Clonetech) was generated by PCR amplification of GCN4-GlyR and subcloned into pmEGFP-C2 using XhoI and KpnI. Full-length gephyrin (isoform P1) G134R was amplified with forward (GACTCAGATCTCGAGCATGGCGACCGAGGGAAT) and reverse (TAGCCGTCCGATGACC) primers and subcloned into pAAV-hSyn-FLEX-mScarlet-Gphn_P1 (Macha et al. 2025) using XhoI and BshTI restriction sites. mEGFP-tagged full-length gephyrin (isoforms P1) was subcloned into pAAV-hSyn-FLEX-mScarlet-Gphn_P1 using NcoI restriction site. The plasmids pAdDeltaF6 (Addgene plasmid # 112867; RRID:Addgene_112867) and pAAV2/1 (Addgene plasmid # 112862; RRID:Addgene_112862) were gifts from James M. Wilson. All DNA constructs were confirmed by Sanger sequencing (Eurofins), and the integrity of AAV2 ITR sites was confirmed by analytical SmaI restriction digest.

### Protein expression, purification and size exclusion chromatography (SEC)

All gephyrin variants used in this study were recombinantly expressed in *E. coli* BL21 (DE3) rosetta cells in 1x LB medium containing 0.1 mg/ml ampicillin. Prior to expression, cells were grown until OD600 of 0.3-0.4 was reached. His-tagged full length gephyrin was expressed for 16 h at 18°C. His-tagged GephG was expressed for 16 h at 30°C. Protein expression was induced addition of isopropyl-β-D-thiogalactopyranosid (IPTG) to an end-concentration 250 μM in the cell-culture. After protein expression, cells were harvested by centrifugation at 4000 g and resuspended in lysis buffer (100 mM Tris/HCl pH 8, 250 mM NaCl, 0.05 % (v/v) Tween20, 5 mM β-mercaptoethanol, 1x protease inhibitor (cOmplete, Roche, Basel, Switzerland), 6.5 mg lysozyme per L expression culture from chicken egg (Sigma-Aldrich) and 25 µg DNAse per mL lysis Buffer) and stored at – 20°C for further use. For purification, the cells were lysed mechanically using EmulsiFlex (Avestin, Ottawa, Canada) and ultrasound for 3x 30 s at 45 % amplitude. Cytosolic extract was separated from cell debris by centrifugation at 50000 g for 45 min. Prior to Ni-NTA chromatography the cell extracts were supplemented with imidazole to 20 mM end concentration. The cell extract was applied to 10 ml Ni-NTA resin and proteins were washed by 10 column volumes (CV) of lysis buffer, 10 CV of lysis buffer containing 10 mM imidazole and 10 CV of lysis buffer containing 20 mM imidazole. Proteins were eluted with 3 CV lysis buffer containing 400 mM imidazole. All proteins were further purified and analyzed via size exclusion chromatography (SEC) (Superdex 16/600 prep grade, GE Healthcare) using protein buffer (20 mM Tris/HCl pH 7.5, 250 mM NaCl, 5 mM β-mercaptoethanol, 5 % glycerol) at a flow rate of 1 mL/min. All purification steps were performed at 4 °C. GlyR-βloop was expressed and purified as stated previously (Grünewald et al. 2018). GlyR-βICD, GABA_A_R-α1ICD and GABA_A_R-α3ICD were purified according to the protocol provided by Bai and colleagues (Bai et al. 2021).

### Isothermal titration calorimetry (ITC)

All experiments were performed with a MicroCal Auto-ITC200 (Malvern, Malvern, United Kingdom) in ITC buffer (20 mM Tris/HCl pH 7.5, 250 mM NaCl, 5 mM β-mercaptoethanol, 5 % glycerol). The sample cell was filled with gephyrin variants at concentration of 20 - 30 μM. GlyR-βloop was filled into the syringe at a concentration of 250-350 μM. Experiments were conducted at 37 °C with an injection volume of 1.5 - 2 μl, a set reference power of 5 μcal/sec, spacing time of 180-300 s and a stirring speed of 750 rpm. Data was analyzed using Origin 7 (OriginLab Corporation) and binding parameters were derived by applying a two-site binding model.

### Sedimentation assay of *in vitro* phase separation

Prior the assay proteins were pre-cleared at 17,000× *g* for 10 min at 25 °C. For sedimentation assays, proteins were mixed at designated combinations and with 10 µM gephyrin and 20 µM receptor model in 50 μL final volume in assay buffer (25 mM Tris pH 7.4, 150 mM NaCl). After 10 min incubation at 25°C, the mixtures were centrifuged at 17,000× *g* for 10 min at 4 °C. Right after centrifugation, the supernatant and pellet were separated. The pellet was brought to the same volume as the supernatant. Proteins recovered in supernatant and pellet were analyzed by SDS-PAGE with Coomassie Blue staining. The band intensities in SDS-PAGE gels were quantified by Image Lab software. Relative and absolute amounts of proteins in supernatant and pellet were calculated based on the relative band intensities. Data of three repeats were presented as means ± SD.

### *In vitro* Moco synthesis assay

The *in vitro* MoCo synthesis assay has previously been described in detail (Belaidi and Schwarz 2013). About 150 pmol *E. coli*-expressed and purified gephyrin was used per condition. Samples without gephyrin or without molybdenum were used as control and show MoCo produced either chemically or by trace amounts of molybdenum in the buffers. Proteins from two independent purifications were used for the assay.

### SDS-PAGE, Coomassie, and Western Blot

Samples were supplemented with 1× sample buffer (5×: 250 mM Tris/HCl pH 6.8, 30% glycerol, 0.1% Bromophenol blue, 10% SDS, 5% β-mercaptoethanol) and incubated for 5 min at 95°C. Protein separation was performed with 12% SDS acrylamide gels, followed by Coomassie staining (30% EtOH, 10% acetic acid, 0.25% Coomassie brilliant blue R250) or immunoblotting using standard protocols with a chemiluminescence and an ECL system. The following antibodies were used and diluted in Tris-buffered saline/0.5% Tween containing 1% dry milk: anti-gephyrin E-domain (3B11, 1:20, self-made, RRID: AB_887719); anti-GAPDH (1:1000, G9545, Sigma), anti-mouse HRP-coupled (1:10 000, AP181P, Sigma); anti-rabbit HRP-coupled (1:10 000, AP187P, Sigma). Image acquisition of Coomassie stained gels and immunoblot detection were performed with a ChemiDocTM Imaging System (BioRad). Band intensities were quantified using Image Lab 6.1 (BioRad).

### Blue Native PAGE Analysis

Samples for blue native PAGE analysis were prepared by 1:3 dilution using H_2_O and the addition of 10× blue native loading buffer (312 mM imidazole, 500 mM 6-aminohexanoic acid, 5% CBB G250, 6.25 mM EDTA, 40% glycerol). Samples were applied to 4%–16% native PAGE (TM Novex Bis-Tris Gel) and protein separation was performed according to the manufacturer’s protocol. Afterward, gels were subjected to western blotting.

### Structural predictions by AlphaFold3

Structural predictions were run in AlphaFold3 (Jumper et al. 2021). For structural comparison the GephG WT (PDB ID: 1JLJ) was used. For G134R substitution the gephyrin WT (UniProt ID: Q9NQX3) sequence was used. Predictions were run three times, with all five output predictions used for analysis. Sequence alignments were conducted on the UniProt website (The UniProt Consortium 2025).

### Generation and transfection of HEK293T *GPHN* KO Cells

Generation of HEK293T *GPHN* KO cells was described by N. Burdina and colleagues (Burdina et al. 2025). 1×10^6^ HEK293T *GPHN* KO cells were plated per well into 6-well cell culture plates and incubated 24 h in Dulbecco’s modified Eagle’s medium supplemented with 10 % FCS and 2 mM l-glutamine at 37 °C and 5% CO_2_. Transfection was performed using 200 ng of the respective plasmid DNA with polyethylenimine (PEI, Sigma) according to the manufacturer’s protocol. 14 h after PEI transfection, cells were washed twice in ice cold PBS and lysed in native lysis buffer (20 mM HEPES pH 8.0, 150 mM NaCl, 5% Glycerol, 5 mM EDTA, 1% DDM, 2x protease inhibitor (cOmplete, Roche, Basel, Switzerland)). Lysates were cleared at 4 °C and 14,000 x g for 20 min and total protein concentration determined using Bicinchoninic Acid assay (Sigma).

### Sulfite oxidase activity in *GPHN* KO HEK293T cells

HEK293T cells (DSMZ no. ACC 635) and *GPHN* KO HEK293T cells were seeded in 6-well plates and co-transfected the next day with human sulfite oxidase in pcDNA5 FRT (Bender et al. 2019) and untagged gephyrin constructs (or empty plasmid, Mock) in pcDNA™3.1/Zeo (+) using Polyethylenimine. 24 h post-transfection cells were washed with PBS and lysed for 1 h at 4 °C (150 mM NaCl, 5 mM EDTA, 1 % (w/v) IGEPAL, 0.5 % (w/v) desoxycholate, 0.1 % (w/v) SDS, 50 mM Tris/HCl pH 7.5 with 1x protease inhibitor mix (cOmplete, Roche, Basel, Switzerland)). Lysates were cleared at 4 °C and 14,000 x g for 20 min and total protein concentration determined using Bicinchoninic Acid assay (Sigma). Sulfite oxidase-specific activity in 100 µg whole cell lysate was measured at room temperature (aerobic conditions) using a plate reader equipped with monochromators (Tecan Spark). In 200 μL, 50 μM cytochrome *c*, 20 μg/mL catalase (bovine, Sigma-Aldrich), 50 µM KCN, and 300 μM Na_2_SO_3_ were combined in 100 mM Tris/Ac pH 8.0 and reduced cytochrome *c* was measured at 550 nm. Initial cytochrome *c* reduction velocities were determined and specific activities calculated using Lambert-Beer Law with ε_550 nm_ = 19 630 M^−1^ cm^−1^ and *d* = 0.857 cm.

### Primary hippocampal cultures

Floxed gephyrin mice (O’Sullivan et al. 2016), a gift from Ralph A. Nawrotzki, were back-crossed to C57BL/6NRj. Mice were housed on a 12 h light-dark cycle with free access to food and water at 22 ± 2 °C and a humidity of 55 ± 10 %. Dissociated primary hippocampal cultures from homozygous gephyrin floxed embryos (E17.5) were prepared of either sex. Embryonal hippocampi obtained from one pregnant mouse were pooled prior to cell dissociation. After dissociation, 65,000 cells were seeded on Poly-L-lysine-coated 13-mm cover slips or 91,000 cells per well on Poly-L-lysine-coated 24-well plates. Cultures were grown in 0.5 mL per well neurobasal medium supplemented with B-27, N-2, GlutaMAX^TM^ (Thermo Fisher Scientific), and Pen/Strep. Recombinant AAV1 particles were diluted in Neurobasal medium with supplements before they were added to the cultures. Cultures were infected (for Western blot analysis) with 1 × 10^8^ viral genome copies (GC) of moxBFP or moxBFP-P2A-Cre or transfected (for immunocytochemistry) with 400 ng DNA using Lipofectamine 2000 after 6 days *in vitro* (DIV) and with 5 × 10^8^ GC of AAV-hSyn-FLEX-mScarlet constructs after after 9 DIV.

### rAAV2/1 preparation

Recombinant AAV2/1 particles were prepared in HEK293T cells (DSMZ no. ACC 635), precipitated with PEG/NaCl, and subsequently cleared via chloroform extraction (Kimura et al. 2019). Purity of preparations was assessed with SDS-PAGE, and titers were determined using Gel green® (Biotium) (Xu et al. 2020).

### Immunocytochemistry

Cells were washed with PBS and fixed at DIV17 with 4 % formaldehyde in PBS for 15 min and quenched with NH_4_Cl in PBS at room temperature. After blocking/permeabilization for 1 h with 10 % goat serum, 1 % BSA, 0.2 % Triton X-100 in PBS, the following primary antibodies (Synaptic Systems) were used: anti-vesicular GABA transporter (vGAT) (1:1000, #131003) for inhibitory presynaptic terminals; anti-GABA_A_R γ2 (1:500, #224004) for postsynaptic GABA_A_Rs. The following secondary antibodies were used: goat anti-rabbit AlexaFluor 488 (1:500, #A-11034, Thermo Fisher Scientific), goat anti-rabbit AlexaFluor 647 (1:500, #A-21245, Thermo Fisher Scientific), and goat anti-guinea pig AlexaFluor 647 (1:500, #ab150187, Abcam). Coverslips were mounted with Mowiol/Dabco.

### Confocal microscopy and image analysis

Image stacks [0.3-μm z-step size, 2520 × 2520 (144.77 × 144.77 μm)] were acquired on a Leica TCS SP8 LIGHTNING upright confocal microscope with an HC PL APO CS2 63×/1.30 glycerol objective, equipped with hybrid detectors (Leica HyD) and the following diode lasers: 405, 488, 552, and 638 nm. LIGHTNING adaptive deconvolution using “Mowiol” setting was applied. Images were segmented and analyzed in an automated fashion using Python 3.10.8 (scikit-image 0.26.0, pyclesperanto-prototype 0.2.1, apoc 0.12.0, numpy 2.3.5, pandas 2.3.3, scipy 1.16.3, napari_simpleitk_image_processing, os, shutil 1.8.1), and the code is available online at GitHub (https://github.com/FilLieb/automated_synapse_analysis).

### Computation

Statistical analyses for image analyses (analysis of variance (ANOVA) with Tukey’s post hoc test, Mann Whitney U test) were performed using R version 4.5.2, rstatix v. 0.7.2, tidyverse v. 2.0.0 in Rstudio environment (https://www.rstudio.com/), Platform: x86_64-w64-mingw32/x64. Data were tested for normality, using violation of the Shapiro–Wilk test at p < 0.01 as the criterion. For all other statistical analyses GraphPad Prism 10 Version 10.2.3 was used for analysis of variance (ANOVA) with Tukey’s multiple comparisons test. Perplexity was used for language style improvement. OMERO, BioRender and ChimeraX-1.8 were used for figure design.

### Ethics statement

We complied with all relevant ethical regulations for animal testing and research and experiments were approved by local ethics committees (Germany, Landesamt für Natur, Umwelt und Verbraucherschutz Nordrhein-Westfalen, reference 2021.A450).

## Acknowledgements

We thank the Biocenter Imaging Facility for their support with confocal microscopy, Dr. Franziska Neuser and Julia Reich, for their support with primary hippocampal cultures, as well as Monika Laurien, and Simona Jansen. We thank PD Dr. med. Ralph A. Nawrotzki, Universität Heidelberg, for kindly sharing floxed gephyrin mice with us. Funding by the German Research Foundation is gratefully acknowledged by GS (RTG2550/1736 project ID 411422114). We also thank Prof. Mingjie Zhang, Shenzhen Southern University of Science and Technology, for kindly sharing their DNA constructs with us.

The authors declare that they have no conflict of interest.

## Notes

### Competing Interest Statement

The authors have declared no competing interest.

## LITERATURE

Bai, Guanhua, Yu Wang, and Mingjie Zhang. 2021. “Gephyrin-Mediated Formation of Inhibitory Postsynaptic Density Sheet via Phase Separation.” Cell Research 31 (3): 3. 10.1038/s41422-020-00433-1.

Bai, Guanhua, and Mingjie Zhang. 2022. “Inhibitory Postsynaptic Density from the Lens of Phase Separation.” Oxford Open Neuroscience 1 (May): kvac003. 10.1093/oons/kvac003.

Bedet, Cécile, Jo C. Bruusgaard, Sandra Vergo, et al. 2006. “Regulation of Gephyrin Assembly and Glycine Receptor Synaptic Stability*.” Journal of Biological Chemistry 281 (40): 30046–56. 10.1074/jbc.M602155200.

Belaidi, Abdel A., and Guenter Schwarz. 2013. “Metal Insertion into the Molybdenum Cofactor: Product–Substrate Channelling Demonstrates the Functional Origin of Domain Fusion in Gephyrin.” Biochemical Journal 450 (1): 149–57. 10.1042/BJ20121078.

Bender, Daniel, Alexander Tobias Kaczmarek, Jose Angel Santamaria-Araujo, et al. 2019. “Impaired Mitochondrial Maturation of Sulfite Oxidase in a Patient with Severe Sulfite Oxidase Deficiency.” Human Molecular Genetics 28 (17): 2885–99. 10.1093/hmg/ddz109.

Burdina, Nele, Filip Liebsch, Arthur Macha, et al. 2025. “Phosphoinositide- and Collybistin-Dependent Synaptic Clustering of Gephyrin.” Journal of Neurochemistry 169 (8): e70169. 10.1111/jnc.70169.

Cross, J. Helen, and Renzo Guerrini. 2013. “Chapter 64 - The Epileptic Encephalopathies.” In Handbook of Clinical Neurology, edited by Olivier Dulac, Maryse Lassonde, and Harvey B. Sarnat, vol. 111. Pediatric Neurology Part I. Elsevier. 10.1016/B978-0-444-52891-9.00064-6.

Dejanovic, Borislav, Tania Djémié, Nora Grünewald, et al. 2015. “Simultaneous Impairment of Neuronal and Metabolic Function of Mutated Gephyrin in a Patient with Epileptic Encephalopathy.” EMBO Molecular Medicine 7 (12): 1580–94. 10.15252/emmm.201505323.

Fritschy, Jean-Marc, Robert J. Harvey, and Günter Schwarz. 2008. “Gephyrin: Where Do We Stand, Where Do We Go?” Trends in Neurosciences 31 (5): 257–64.

Grünewald, Nora, Audric Jan, Charlotte Salvatico, et al. 2018. “Sequences Flanking the Gephyrin-Binding Site of GlyRβ Tune Receptor Stabilization at Synapses.” New Research. eNeuro 5 (1). 10.1523/ENEURO.0042-17.2018.

Guerrini, Renzo, Valerio Conti, Massimo Mantegazza, Simona Balestrini, Aristea S. Galanopoulou, and Fabio Benfenati. 2023. “Developmental and Epileptic Encephalopathies: From Genetic Heterogeneity to Phenotypic Continuum.” Physiological Reviews 103 (1): 433–513. 10.1152/physrev.00063.2021.

Herweg, Jens, and Guenter Schwarz. 2012. “Splice-Specific Glycine Receptor Binding, Folding, and Phosphorylation of the Scaffolding Protein Gephyrin.” The Journal of Biological Chemistry 287 (16): 12645–56. 10.1074/jbc.M112.341826.

Johannes, Lena, Chun-Yu Fu, Günter Schwarz, Lena Johannes, Chun-Yu Fu, and Günter Schwarz. 2022. “Molybdenum Cofactor Deficiency in Humans.” Molecules 27 (20). 10.3390/molecules27206896.

Jumper, John, Richard Evans, Alexander Pritzel, et al. 2021. “Highly Accurate Protein Structure Prediction with AlphaFold.” Nature 596 (7873): 583–89. 10.1038/s41586-021-03819-2.

Kim, Eun Young, Nils Schrader, Birthe Smolinsky, et al. 2006. “Deciphering the Structural Framework of Glycine Receptor Anchoring by Gephyrin.” The EMBO Journal 25 (6): 1385–95. 10.1038/sj.emboj.7601029.

Kim, Seungjoon, Mooseok Kang, Dongseok Park, et al. 2021. “Impaired Formation of High-Order Gephyrin Oligomers Underlies Gephyrin Dysfunction-Associated Pathologies.” iScience 24 (2). 10.1016/j.isci.2021.102037.

Kimura, Toyokazu, Beatriz Ferran, Yuko Tsukahara, et al. 2019. “Production of Adeno-Associated Virus Vectors for in Vitro and in Vivo Applications.” Scientific Reports 9 (1): 13601. 10.1038/s41598-019-49624-w.

Lee, Gyehyun, Seungjoon Kim, Da-Eun Hwang, et al. 2024. “Thermodynamic Modulation of Gephyrin Condensation by Inhibitory Synapse Components.” Proceedings of the National Academy of Sciences 121 (12): e2313236121. 10.1073/pnas.2313236121.

Liebsch, Filip, Fynn R. Eggersmann, Yvonne Merkler, Peter Kloppenburg, and Günter Schwarz. 2023. “Automated Image Analysis Reveals Different Localization of Synaptic Gephyrin C4 Splice Variants.” Research Article: New Research. eNeuro 10 (1). 10.1523/ENEURO.0102-22.2022.

Lionel, Anath C., Andrea K. Vaags, Daisuke Sato, et al. 2013. “Rare Exonic Deletions Implicate the Synaptic Organizer Gephyrin (GPHN) in Risk for Autism, Schizophrenia and Seizures.” Human Molecular Genetics 22 (10): 2055–66. 10.1093/hmg/ddt056.

Macha, Arthur, Filip Liebsch, Nele Burdina, et al. 2025. “Gephyrin Filaments Represent the Molecular Basis of Inhibitory Postsynaptic Densities.” Preprint, Research Square, March 13. 10.21203/rs.3.rs-5889959/v1.

Macha, Arthur, Filip Liebsch, Steffen Fricke, et al. 2022. “Biallelic Gephyrin Variants Lead to Impaired GABAergic Inhibition in a Patient with Developmental and Epileptic Encephalopathy.” Human Molecular Genetics 31 (6): 901–13. 10.1093/hmg/ddab298.

Maric, Hans Michael, Vikram Babu Kasaragod, Torben Johann Hausrat, et al. 2014. “Molecular Basis of the Alternative Recruitment of GABAA versus Glycine Receptors through Gephyrin.” Nature Communications 5 (1): 5767. 10.1038/ncomms6767.

Maric, Hans-Michael, Jayanta Mukherjee, Verena Tretter, Stephen J. Moss, and Hermann Schindelin. 2011. “Gephyrin-Mediated γ-Aminobutyric Acid Type A and Glycine Receptor Clustering Relies on a Common Binding Site*.” Journal of Biological Chemistry 286 (49): 42105–14. 10.1074/jbc.M111.303412.

Meyer, G., J. Kirsch, H. Betz, and D. Langosch. 1995. “Identification of a Gephyrin Binding Motif on the Glycine Receptor Beta Subunit.” Neuron 15 (3): 563–72. 10.1016/0896-6273(95)90145-0.

Mittag, Tanja, and Rohit V. Pappu. 2022. “A Conceptual Framework for Understanding Phase Separation and Addressing Open Questions and Challenges.” Molecular Cell 82 (12): 2201–14. 10.1016/j.molcel.2022.05.018.

O’Sullivan, Gregory A., Peter Jedlicka, Hong-Xing Chen, et al. 2016. “Forebrain-Specific Loss of Synaptic GABAA Receptors Results in Altered Neuronal Excitability and Synaptic Plasticity in Mice.” Molecular and Cellular Neurosciences 72 (April): 101–13. 10.1016/j.mcn.2016.01.010.

Reinthaler, Eva M., Borislav Dejanovic, Dennis Lal, et al. 2015. “Rare Variants in γ-Aminobutyric Acid Type A Receptor Genes in Rolandic Epilepsy and Related Syndromes.” Annals of Neurology 77 (6): 972–86. 10.1002/ana.24395.

Saiyed, Taslimarif, Ingo Paarmann, Bertram Schmitt, et al. 2007. “Molecular Basis of Gephyrin Clustering at Inhibitory Synapses: Role of G-and E-Domain Interactions.” Journal of Biological Chemistry 282 (8): 5625–32.

Sander, Bodo, Giancarlo Tria, Alexander V. Shkumatov, et al. 2013. “Structural Characterization of Gephyrin by AFM and SAXS Reveals a Mixture of Compact and Extended States.” Acta Crystallographica. Section D, Biological Crystallography 69 (Pt 10): 2050–60. 10.1107/S0907444913018714.

Scheffer, Ingrid E., Jacqueline French, Kette D. Valente, Stéphane Auvin, J. Helen Cross, and Nicola Specchio. 2025. “Operational Definition of Developmental and Epileptic Encephalopathies to Underpin the Design of Therapeutic Trials.” Epilepsia 66 (4): 1014–23. 10.1111/epi.18265.

Schwartzkroin, Philip A. 2012. “Chapter 2 - Cellular Bases of Focal and Generalized Epilepsies.” In Handbook of Clinical Neurology, edited by Hermann Stefan and William H. Theodore, vol. 107. Epilepsy. Elsevier. 10.1016/B978-0-444-52898-8.00002-1.

Schwarz, G., N. Schrader, R. R. Mendel, H. J. Hecht, and H. Schindelin. 2001. “Crystal Structures of Human Gephyrin and Plant Cnx1 G Domains: Comparative Analysis and Functional Implications.” Journal of Molecular Biology 312 (2): 405–18. 10.1006/jmbi.2001.4952.

Schwarz, Günter, Ralf R. Mendel, and Markus W. Ribbe. 2009. “Molybdenum Cofactors, Enzymes and Pathways.” Nature 460 (7257): 839–47. 10.1038/nature08302.

Shen, Zeyu, Bowen Jia, Yang Xu, et al. 2023. “Biological Condensates Form Percolated Networks with Molecular Motion Properties Distinctly Different from Dilute Solutions.” eLife 12 (June): e81907. 10.7554/eLife.81907.

Tyagarajan, Shiva K., and Jean-Marc Fritschy. 2014. “Gephyrin: A Master Regulator of Neuronal Function?” Nature Reviews Neuroscience 15 (3): 141–56.

Uzay, Burak, and Ege T. Kavalali. 2023. “Genetic Disorders of Neurotransmitter Release Machinery.” Frontiers in Synaptic Neuroscience 15 (March): 1148957. 10.3389/fnsyn.2023.1148957.

Xu, Jian, Steven H. DeVries, and Yongling Zhu. 2020. “Quantification of Adeno-Associated Virus with Safe Nucleic Acid Dyes.” Human Gene Therapy 31 (19–20): 1086–99. 10.1089/hum.2020.063.

Yardi, Ruta. 2025. “Portrait of Epilepsy on the Canvas of Global Health.” Epilepsy Currents, June 25, 15357597251352422. 10.1177/15357597251352422.

Zhu, Shihan, Zeyu Shen, Xiandeng Wu, et al. 2024. “Demixing Is a Default Process for Biological Condensates Formed via Phase Separation.” Science 384 (6698): 920–28. 10.1126/science.adj7066.

